# N-terminal intrinsically disordered region mediates self-catalytic interfacial nucleation of *Aspergillus oryzae* hydrophobin RolA

**DOI:** 10.64898/2026.08.04.742928

**Authors:** Nao Takahashi, Natsuki Abe, Takuya Mabuchi, Mao Fukuyama, Yuki Terauchi, Takumi Tanaka, Akira Yoshimi, Hiroshi Yabu, Keietsu Abe

## Abstract

Hydrophobins are biosurfactant proteins that coat the cell surfaces of filamentous fungi. On the conidial surface, hydrophobins self-assemble into rodlets, forming a dense hydrophobic film that promotes air-dispersibility. Although rodlet formation is closely associated with the physiology of filamentous fungi, its underlying molecular mechanisms remain largely unknown. Previously, we revealed that RolA, a hydrophobin derived from *Aspergillus oryzae*, forms rodlets at the air–water interface. In this study, we focused on the flexible N-terminal region of RolA, which lacks a well-defined tertiary structure, and hypothesized that this intrinsically disordered region regulates rodlet formation. To investigate its role, we used RolA mutants with reduced charges in the N-terminal region and analyzed the rodlet formation process on the surface of a water-in-air sessile droplet using atomic force microscopy. In addition, we quantitatively characterized rodlet formation at the air–water interface by applying a kinetic perspective to the interfacial tension change profiles obtained from dynamic surface tension measurements. The results suggested that RolA first forms a monolayer at the air–water interface, then rodlet formation proceeds through the continuous supply of free RolA monomers from the bulk phase to the interfacial RolA film. Our molecular dynamics simulations of RolA at the interface supported a model in which RolA molecules within the interfacial film interact with free monomers in the bulk phase through their N-terminal regions. These results reveal a previously unidentified role of the N-terminal region in rodlet formation and provide a more comprehensive framework for understanding the molecular mechanism underlying RolA rodlet formation.

## INTRODUCTION

Amyloid proteins undergo self-assembly to form fibrous structures. Although these structures are typically associated with neurodegenerative diseases, functional amyloids contributing to essential biological processes such as morphogenesis have also been identified [1,2]. Amyloids are attracting increasing attention not only as targets for drug discovery but also as building blocks for protein-based functional materials [3]. Hydrophobin, secreted by filamentous fungi, is recognized as a kind of functional amyloid because it forms rod-shaped multimeric structures called rodlets, which are composed of cross- β-sheet structures similar to those of pathogenic amyloid fibrils, and its role in filamentous fungal physiology has been studied for over two decades. Hydrophobins are localized on the fungal cell surface, making the hydrophilic cell wall hydrophobic, which promotes adhesion of hyphae to solid polymeric substrates and aerial dispersion of conidia [4,5]. Hydrophobins also possess immune-evasive properties, which enable pathogenic filamentous fungi to avoid recognition by the host immune system and thereby contribute to infection [6,7]. In addition, hydrophobins enhance polymer degradation by covering solid polymer surfaces and recruiting and concentrating polymer-degrading enzymes to the polymer surface [8–11]. Owing to these diverse functionalities, hydrophobins have been explored for applications such as drug delivery, emulsification, and the promotion of plastic degradation [12,13].

Hydrophobins generally contain eight cysteine residues in their amino acid sequences, and their 3D structure is stabilized by the four disulfides bonds between these residues [14]. Monomeric hydrophobin possesses a central β-barrel core region while exposing three hydrophobic loop regions (Cys3–Cys4, Cys4–Cys5, Cys7–Cys8) on the molecular surface, and at least one of the three loops has been identified as amyloidogenic [7,15]. Hydrophobins form rodlets through zipper-like stacking of the three hydrophobic loops [15,16]. Rodlets are extremely stable and dissociate only when exposed to strong acids such as trifluoroacetic acid or formic acid [17,18]. Because disruption of the tertiary structure of the monomer interrupts the rodlet-forming ability, proper folding of the monomer is considered essential for rodlet formation [7].

Rodlet formation is strongly promoted at hydrophobic–hydrophilic interfaces such as the air–water interface [10,19–21]. For several hydrophobins, it has been reported that rodlets do not form in bulk solution; instead, an amorphous monolayer is first formed at the interface, followed by rodlet formation [7,21,22]. It is beginning to appear that these two structural states at the interface express distinct functionalities. In our previous study, we demonstrated that the hydrophobin RolA from *Aspergillus oryzae*, an industrial fungus, likely adopts the rodlet state to form an elastic, surface-active interfacial film and make the conidial surface hydrophobic [20]. Thus, understanding the functionality of RolA requires going beyond the monomeric state, and the emerging view that self-assembly governs functional regulation further highlights the physicochemical and biological importance of rodlet formation. However, the rodlet formation mechanism, which plays a central role in the new paradigm, remains largely elusive. Elucidating the rodlet formation mechanism of RolA would enable a better understanding of the diverse functional mechanisms of RolA that are mediated by such a dynamic structural transition.

To reveal the rodlet formation mechanism, we considered it useful to focus on the function of the N-terminal region of RolA. In hydrophobins, the role of the N-terminal region remains far less well understood than that of the loop regions, which are considered crucial for rodlet formation. Although the N-terminal region in several hydrophobins has not been thought to contribute to rodlet formation [7,23], it cannot be ruled out that the insufficient exploration of N-terminal function has obscured the true nature of rodlet formation. In RolA, N-terminal region constitute an intrinsically disordered region (IDR) enriched in charged amino acid residues [9] (see Figure 1). His32 and Lys34 in the N-terminus have already been identified as contributing to the strong interaction between RolA and the hydrolytic enzyme CutL1 via electrostatic interactions [9,11]. IDRs generally lack a single well-defined folded structure and instead behave as a diverse conformational ensemble that adapts to the surrounding environment and binding partners, enabling multivalent and reversible interactions, and functioning as hubs in a wide range of biological process [25]. Therefore, we hypothesized that the RolA N-terminus, by making use of its conformational flexibility, may serve as a bifunctional segment that mediates both CutL1 recruitment and rodlet formation. Indeed, both the RolA–CutL1 interaction and rodlet formation are regulated by electrostatic interactions [9,19].

**Figure 1.**
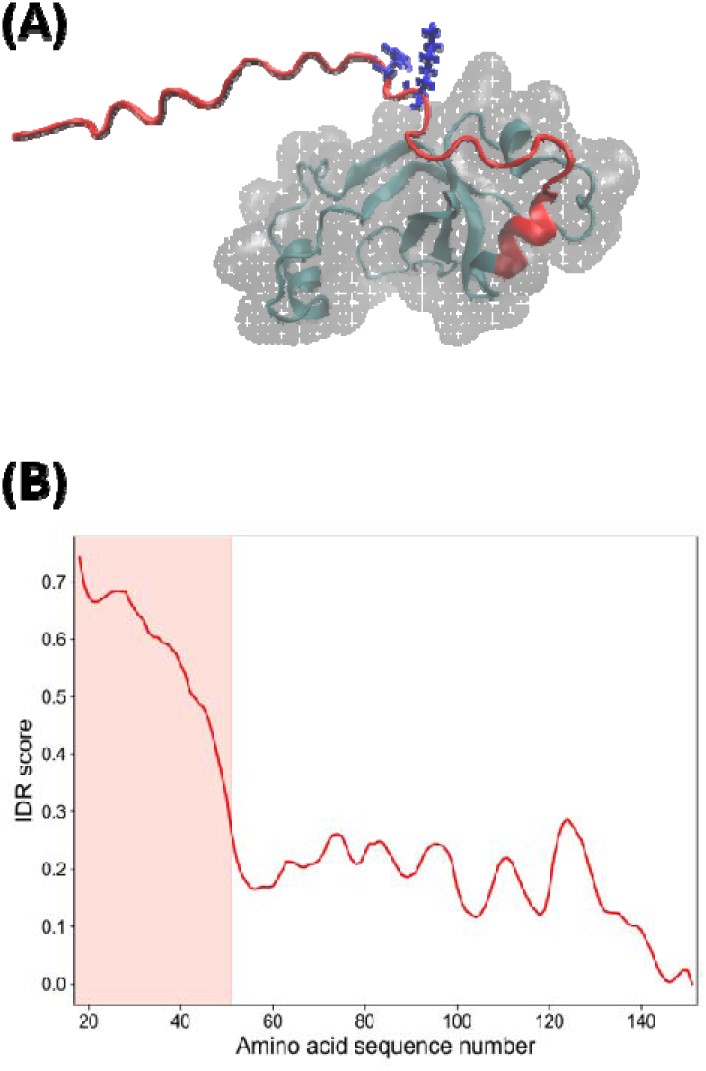
Structural features of RolA. (A) AlphaFold model, in which the N-terminal region is colored red and His32 and Lys34 are shown as blue sticks. (B) intrinsic disorder propensity (IDR) of each residue in RolA predicted using IUPred3 [24].

In this study, we aimed to elucidate how positively charged residues within the RolA N-terminus affect rodlet formation to test the possibility that the RolA N-terminus is bifunctional. Because RolA rodlet formation proceeds only at hydrophobic–hydrophilic interfaces [26], we evaluated rodlet formation kinetics on a droplet surface by atomic force microscopy (AFM). We identified that rodlet formation was delayed in mutants with reduced charge in the N-terminus. Then, to analyze the effects of the mutations in detail, we conducted dynamic surface tension measurements to separately quantify adsorption of RolA to the interface and subsequent rodlet formation. These measurements revealed that RolA first forms an amorphous film at the interface, after which rodlet formation occurs on the surface of the aqueous-phase side of this interfacial film. The results further suggest that the N-terminus specifically modulates nucleation processes beneath the interfacial film, and this model is supported by our molecular dynamics (MD) simulations. These findings update the current model of the RolA rodlet formation process by revealing a mechanism mediated by the RolA N-terminus, highlighting the functional versatility of the RolA N-terminus.

## RESULTS

### Rodlet formation kinetics on droplet surfaces and their structure

To identify the rodlet formation tendency of RolA-WT and three charge-reduced N-terminal mutants (RolA-H32S, -K34S, and -H32S/K34S), we evaluated the rodlet formation speed at the air–water interface of sessile water droplets using the method described by Takahashi *et al*. [19] with slight modifications (Figure 2A). The results show that the coverage ratio of the sessile water droplet surface with rodlets increased with increasing incubation time for RolA-WT and all variants, eventually reaching saturation (Figure 2B). Rodlet formation tended to be suppressed in the RolA variants relative to RolA- WT, with a significant delay in the half-time (the time required to reach 50% coverage) observed particularly for RolA- H32S and -H32S/K34S (Figure 2C).

**Figure 2.**
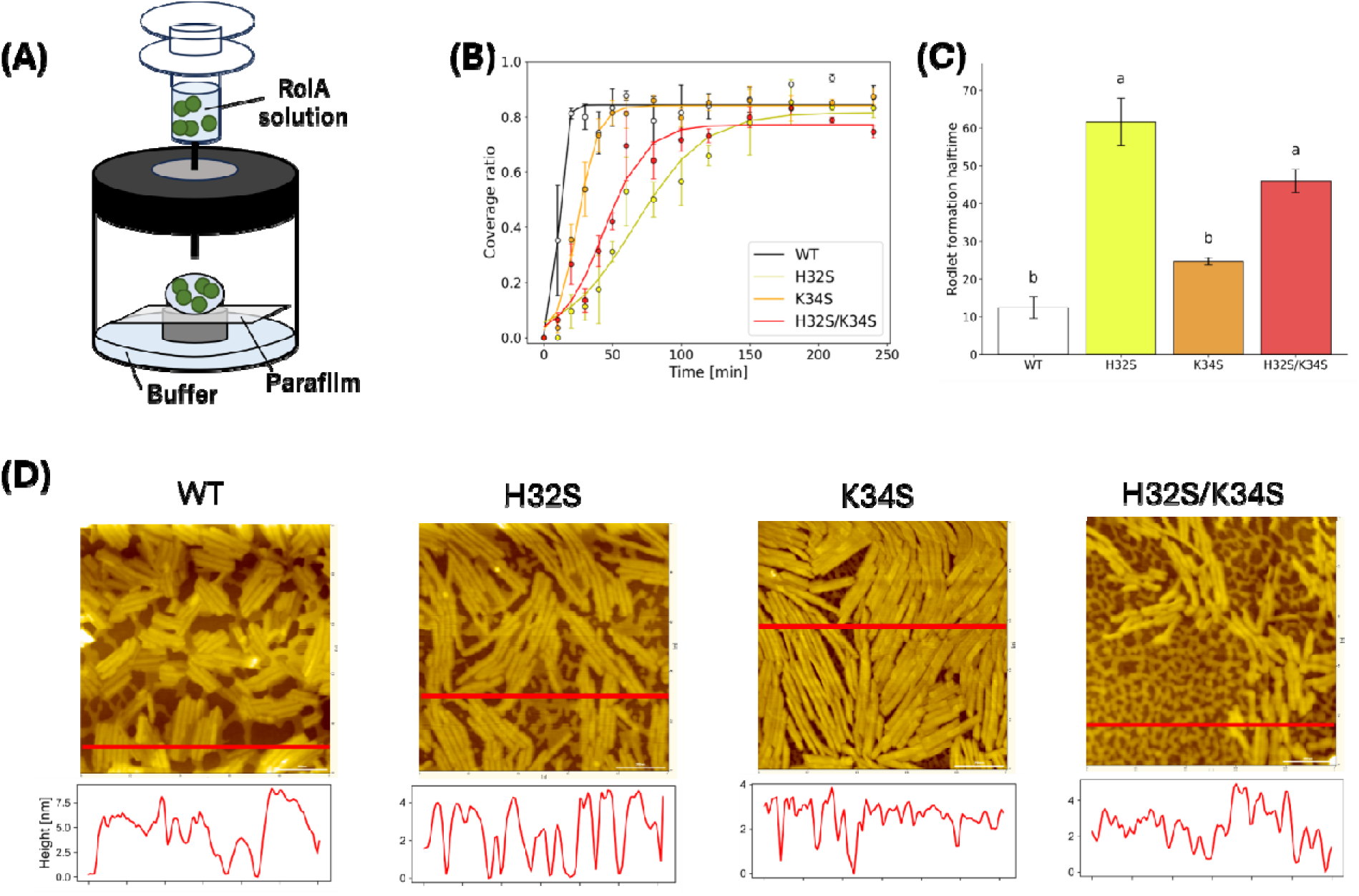
Rodlet formation of RolA-WT and mutants. (A) Schematic illustration of the preparation of a sessile water droplet in air. RolA self-assembles into rodlets at the air–water interface of the droplet. (B) Time course of rodlet surface coverage on the droplet surface. Error bars represent standard error. (C) Half-time required to reach saturation. Error bars represent standard error. Categories with different letters are significantly different at *P* < 0.05. Statistical significance was evaluated using Tukey’s method. (D) Atomic force microscopy image of the intermediate stage before surface saturation. Scale bar, 200 nm. The images shown for WT, K34S, and H32S/K34S were acquired at 40 min, and the image for H32S was acquired at 80 min. Cross-sectional plots along the red lines are shown below each image.

AFM images acquired at the initial stage of rodlet film formation showed that, in addition to rodlets, amorphous-like structures were also present. These structures were lower in height than the rodlets, indicating that RolA forms a multicomponent interfacial film composed of rodlets and other non-rodlet structures (Figure 2D). The distribution of rodlet length extracted from AFM images after 240 min of incubation showed that the mutants tended to form longer rodlets than the WT (Figure S1).

### Surface adsorption and rodlet formation speed revealed by dynamic surface tension measurements

Dynamic surface tension measurements of RolA have shown a two-step decrease in surface tension; the steps correspond to (□) surface adsorption and (□) rodlet formation [20]. Therefore, we sought to quantitatively distinguish and evaluate surface adsorption and rodlet formation by kinetically interpreting this two-step surface tension reduction.

Figure 3A shows the results of dynamic surface tension measurements when the initial protein concentration was varied from 0.78 to 250 μg/mL. At higher initial protein concentration (12.5–250 μg/mL), the surface tension first decreased rapidly to approximately 55 mN/m, followed by an equilibrium phase, and then decreased again to approximately 35 mN/m. At lower concentrations (3.13, 6.25 μg/mL), it took a long time for the surface tension to decrease and reach the first equilibrium, and a clear secondary decrease was not observed. When the concentration was further reduced (0.78 and 1.56 μg/mL), even the primary decrease was not observed. As the initial protein concentration increased, the half-time of the primary decrease in surface tension became shorter (Figure 3B), and the duration of the first equilibrium plateau (lag time) was also shortened (Figure 3C).

**Figure 3.**
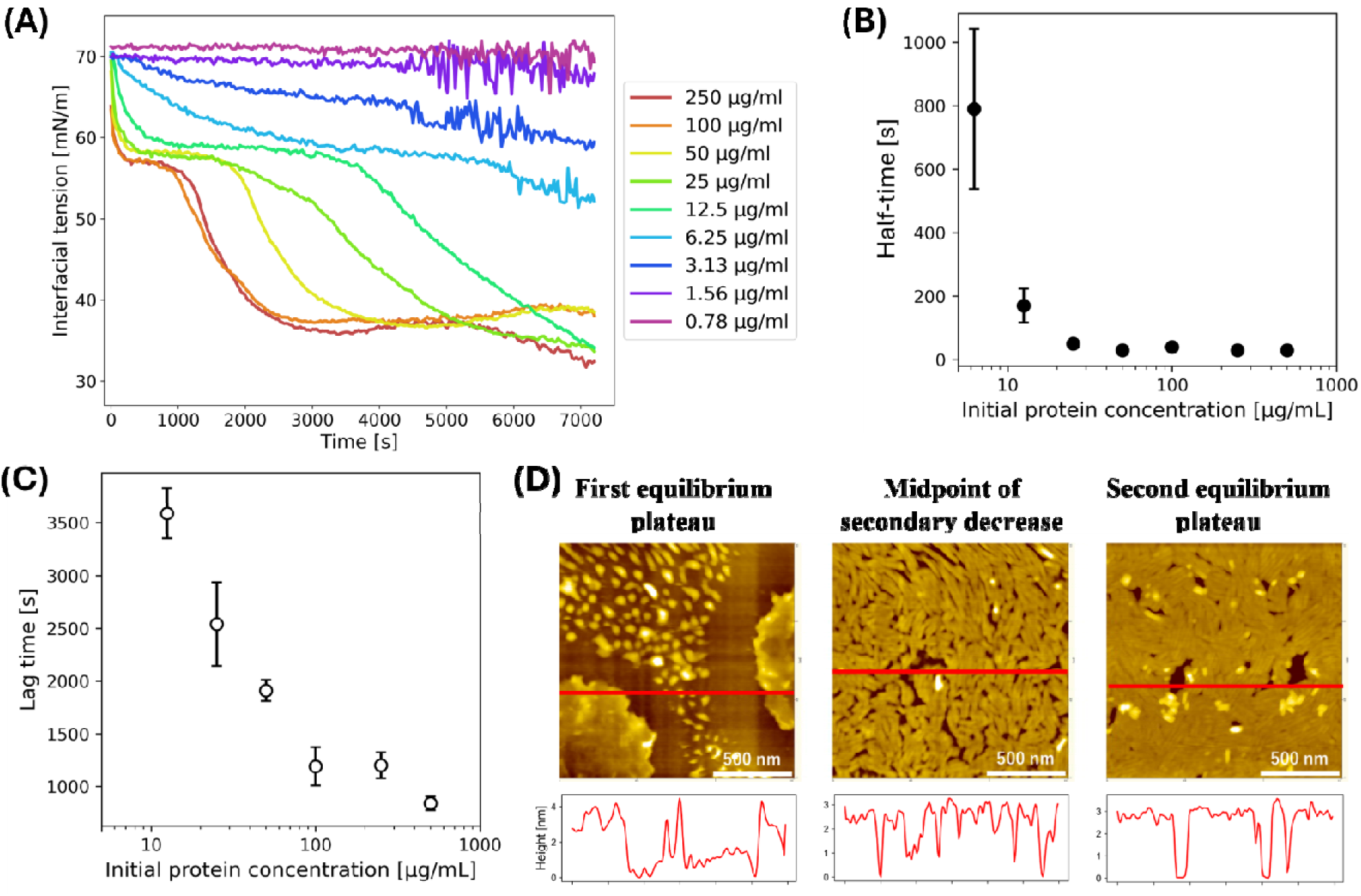
Dynamic surface tension of RolA-WT. (A) Time course of surface tension changes in RolA-WT at different initial protein concentrations. (B) Initial protein concentration dependence of the half-time required to reach the first equilibrium state of surface tension. Error bars represent the standard error. (C) Initial protein concentration dependence of the lag time during which the surface tension remained at the first equilibrium state. Error bars represent standard error. (D) Atomic force microscopy images obtained during dynamic surface tension measurements at 100 μg/mL: at the first equilibrium state, during the second decrease in surface tension, and at the second equilibrium state. Scale bars, 500 nm. Cross-sectional plots along the red lines are shown below each image.

To analyze the RolA structures formed at the droplet surface during the dynamic surface tension measurement, we obtained interfacial structures at three points: (□) the first equilibrium plateau (approximately 55 mN/m), (□) the midpoint of the secondary decrease (approximately 45 mN/m), and (□) the second equilibrium plateau (approximately 35 mN/m) (Figure 3D). AFM analysis showed that rodlets were not observed at the first equilibrium plateau; instead, spherical structures were observed. Once the second decrease began, many rodlets were observed, and at the second equilibrium plateau (the endpoint of the reaction) a dense rodlet film was formed, indicating that the interfacial changes up to the first equilibrium plateau reflect RolA adsorption to the interface, whereas the subsequent process reflects an increase in the amount of rodlet formation at the interface.

### Effect of mutations in the N-terminal region

To assess the effects of mutations in the RolA N-terminal region, dynamic surface tension measurements for the RolA mutants (RolA-H32S, K34S, and H32S/K34S) were also performed (Figure S2). The half-time of the primary decrease in surface tension was significantly longer for RolA-H32S and -H32S/K34S than for -WT and -K34S (Figure 4A), indicating slow surface adsorption for RolA-H32S and -H32S/K34S. When 150 mN of NaCl was added to inhibit electrostatic interactions between RolA molecules, the half-times for RolA-H32S and -H32S/K34S were markedly shortened with the addition of NaCl, resulting in similar half-times among RolA-WT and all mutants (Figure 4B).

**Figure 4.**
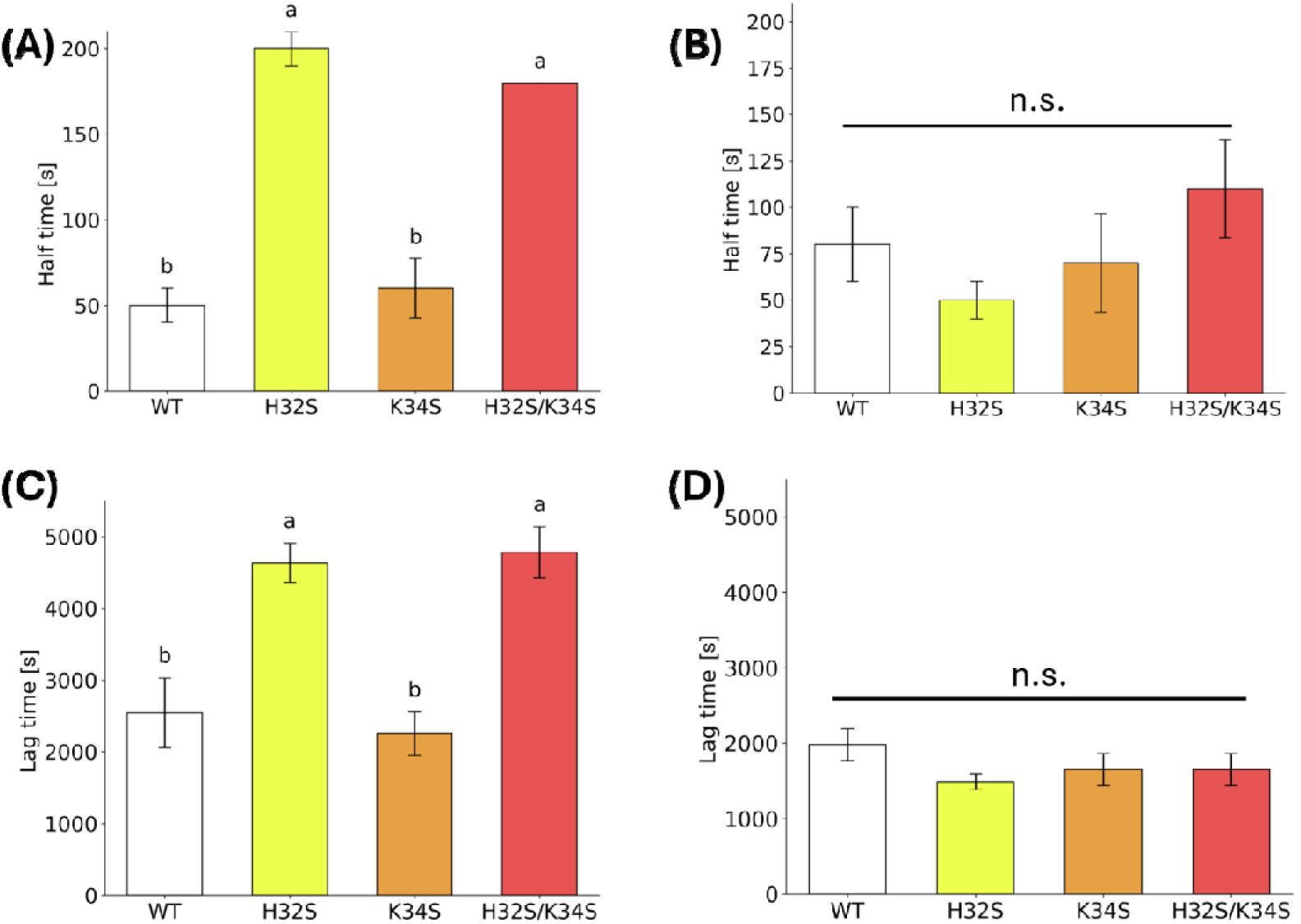
Comparison of half-time and lag time values derived from dynamic surface tension measurements at 25 μg/mL for RolA-WT and mutants. Half-times for reaching the first equilibrium state of surface tension in (A) the absence of NaCl and (B) the presence of 150 mM NaCl. Lag times (where the surface tension remained at the first equilibrium state) in (C) the absence of NaCl and (D) the presence of 150 mM NaCl. Error bars indicate standard error. Categories with different letters are significantly different at *P* < 0.05. Statistical significance was evaluated using Tukey’s method. N.S., not significant.

The lag time was longer for RolA-H32S and RolA-H32S/K34S (Figure 4C), indicating delayed rodlet formation in these mutants. The addition of 150 mM of NaCl shortened the lag time for RolA-H32S and -H32S/K34S, resulting in the disappearance of the differences in lag time among RolA-WT and all mutants (Figure 4D).

The surface tension at the first equilibrium plateau was then plotted as a function of the initial protein concentration (Figure S3). Using Eq. 1, the maximum surface excess concentration (*Γ*) was determined, and the critical micelle concentration (CMC) was estimated (Table S1). *Γ* and CMC varied among the WT and mutants, further supporting the conclusion that mutagenesis can modulate the surface adsorption and ordering of interfacial RolA molecules.

### MD simulations of RolA at the air–water interface

To elucidate the molecular mechanisms by which the N-terminal region of RolA affects rodlet formation, MD simulations were performed. An air–water interface was introduced into the system, and the interfacial structures of RolA monomers were simulated (Figure 5A). The simulations showed that, after adsorption to the interface, RolA adopted an orientation in which the Cys4–Cys5 loop faced the air phase, whereas the Cys3–Cys4 loop and the N-terminal region were positioned toward the aqueous phase. The Cys7–Cys8 loop occupied an intermediate position (Figure 5B).

**Figure 5.**
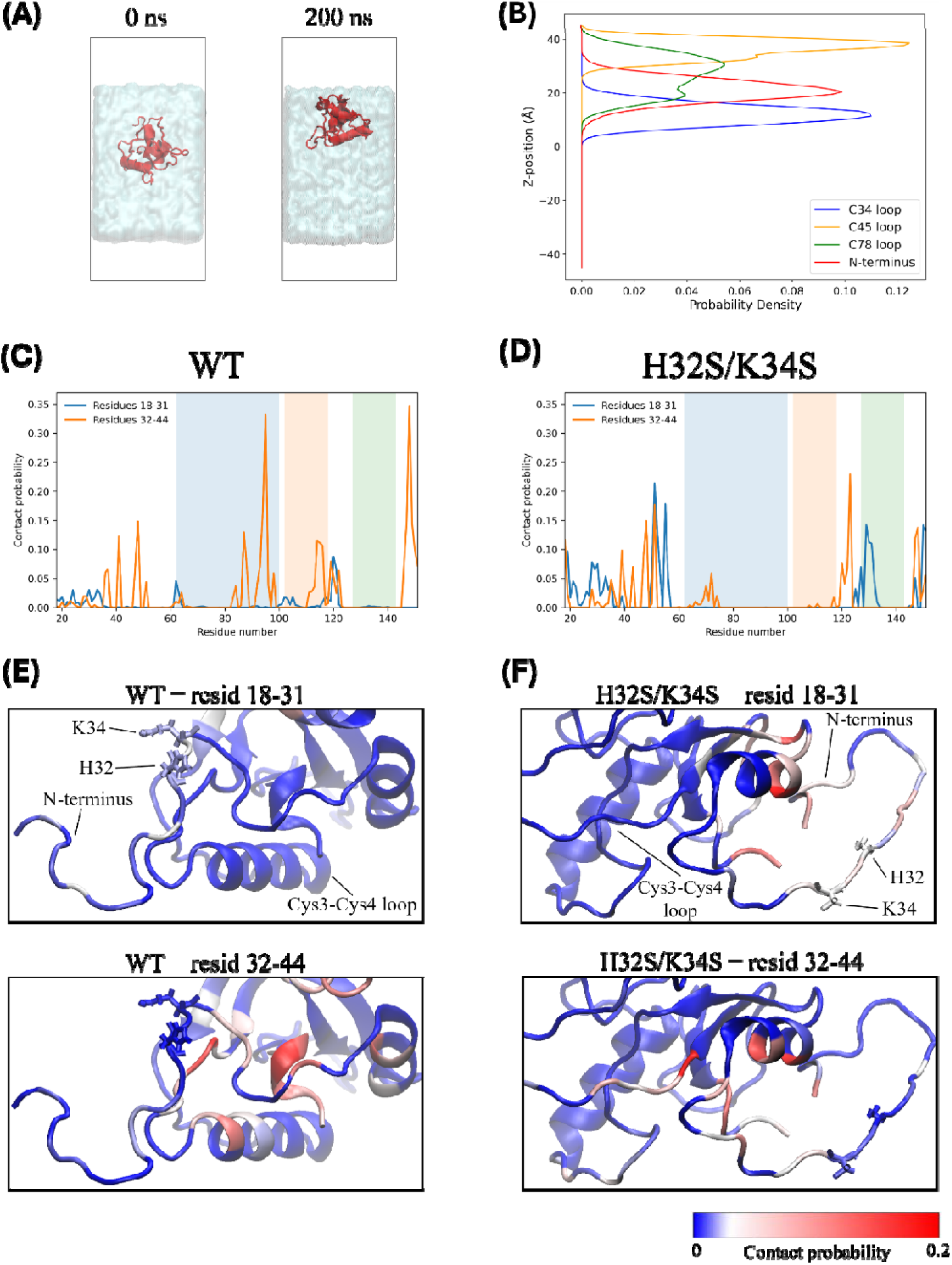
Molecular dynamics simulation of the structure and orientation of RolA at the air–water interface. (A) Snapshots of the adsorption process of RolA at the air–water interface. (B) Spatial distribution of the Cys loops and the N-terminal region. Contact probability profile of (C) WT and (D) H32S/K34S. Contacts between the N-terminal region (comprising residues 18–31) and other regions are shown by blue lines; those between residues 32–44 and other regions are shown by orange lines. The Cys3–Cys4, Cys4–Cys5, and Cys7–Cys8 loops are indicated by blue, orange, and green shading, respectively. Representative structures of interfacial (E) RolA-WT and (F) -H32S/K34S. In the upper panels, colors indicate average contact probabilities between residues 18–31 and other regions. In the lower panels, colors indicate average contact

To compare the structural dynamics of RolA-WT and -H32S/K34S at the interface, root-mean-square fluctuation (RMSF) values were calculated for both variants (Figure S4). H32S/K34S showed increased fluctuation in the Cys3–Cys4 loop relative to the WT. By contrast, within the N-terminal region, fluctuations were substantially reduced in the first half of the region, while those in the latter half remained similar to the WT.

Then, to elucidate the structural state of the N-terminal region at the interface, we analyzed its contacts with other regions of the RolA molecule. Because the RMSF analysis revealed distinct dynamic behaviors between the first and second halves of the N-terminal region, and because the latter half has a distinctive sequence feature characterized by a concentration of charged amino acid residues, the N-terminal region was divided into two segments, residues 18–31 (LPPASGTGAGQQVG) and 32–44 (HSKNDFPLPKELT), and their contact patterns were analyzed separately. In RolA- WT, the first N-terminal segment exhibited few contacts with other regions, whereas the second segment showed a tendency to form transient contacts with the latter part of the Cys3–Cys4 loop (residues 92–100, SQGLGLLDE) and the C-terminus (Figure 5C). In RolA-H32S/K34S, compared with WT, the first N-terminal segment exhibited increased intramolecular contacts within the N-terminal region, whereas contacts between the second N-terminal segment and the Cys3–Cys4 loop were almost lost (Figure 5D). These results indicate that the mutations reduced the tendency of the N-terminal region to approach the Cys3–Cys4 loop and instead promoted intramolecular interactions within the N-terminal region, thereby conformationally constraining its first segment (Figure 5E, F).

## DISCUSSION

In this study, we investigated how the N-terminal IDR of RolA affects rodlet formation. Because RolA rodlet formation is an interface-specific process, quantitative kinetic analyses were performed through AFM observation of interfacial structures and dynamic surface tension measurements. The molecular model suggested by the experimental results was further supported by MD simulations.

AFM-based analysis of rodlet formation at the air–water interface of a sessile water droplet revealed that reducing the positive charges within the N-terminal region suppresses rodlet formation (Figure 2B, C). Previously, the N-terminal region of hydrophobins has been considered not to have significant effects on rodlet formation [7,23]. One reason for this might be that the rodlet formation process has not been clearly modeled, making it difficult to identify which step in the reaction pathway is promoted or inhibited in response to the environmental conditions and mutations. The aggregation pathway of amyloid proteins is usually assessed by thioflavin T (ThT) assay, and this method has also been applied to the evaluation of the rodlet formation mechanism of hydrophobins [10,23,26]. However, a kinetic study of rodlet formation by MPG1, a class-I hydrophobin, showed that the ThT assay cannot distinguish between monomer adsorption to the interface and dock– lock-type rodlet elongation [10], meaning that the strong interface dependence of rodlet formation may obscure the underlying mechanism of rodlet formation in a ThT assay. In this study, we tracked rodlet formation on a sessile water droplet surface and found that rodlet formation was slower in RolA N-terminal mutants (Figure 2B, C). Differences between the WT and mutants (Figure S1) were further evident in the distribution of rodlet lengths. When elongation dominates over nucleation, fewer nuclei are generated, allowing the available protein mass to be incorporated into a small number of growing assemblies and thereby yielding longer fibrils [27]. Accordingly, the longer rodlets observed for the mutants are consistent with reduced nucleation compared with that of WT. Since the isoelectric points of the mutants (H32S, 4.73; K34S, 4.72; H32S/K34S, 4.55) were shifted to the acidic side relative to that of the WT (4.93), the mutants were likely more highly charged at pH 5, leading to stronger electrostatic repulsion and delayed rodlet formation. However, this assay still could not distinguish surface adsorption from rodlet formation and therefore did not essentially resolve the limitation of previous approaches. Consequently, it remained difficult to determine in detail which step was affected by the mutations and to what extent.

To address this issue, we conducted dynamic surface tension measurements using the pendant-drop method. The results showed that RolA-WT (25 μg/mL) reached adsorption equilibrium within approximately 50 s (Figure 3B), whereas approximately 2500 s elapsed before the onset of the secondary decrease in interfacial tension attributable to rodlet formation (Figure 3C). These results indicate that surface adsorption is approximately two orders of magnitude faster than rodlet formation for RolA, suggesting that surface adsorption is unlikely to be rate-limiting for the entire reaction process. Instead, rodlet formation at the interface should be the rate-limiting step.

Because the duration of the first equilibrium plateau corresponds to the waiting time required for sufficient rodlet formation to induce a further decrease in surface tension, this period can be interpreted as a macroscopic proxy for nucleation (lag time). Accordingly, the lag time provides important insight into the interface-specific kinetics of rodlet formation. In kinetic analyses of amyloid fibril formation, characteristic times such as the lag time commonly exhibit a power-law dependence on the initial protein concentration, 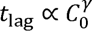, where the scaling exponent is related to the reaction orders and relative contributions of the elementary steps governing the overall assembly process [28]. Based on this general scaling behavior, the initial concentration dependence of the lag time observed in the present system was fitted using the following equation:

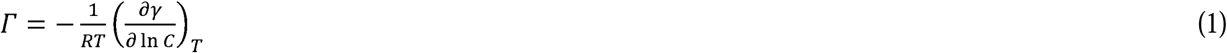

where *A* is a constant, *C*_0_ is the initial bulk protein concentration, γ is the scaling factor, and *t*_0_ is a concentration-independent offset. When the surface tension reached the first equilibrium plateau, the air–water interface was considered to be saturated with adsorbed RolA molecules, such that the interface concentration reached an approximately constant surface excess, here donated as *C*_eq_ (Table S1). Given that RolA forms rodlets specifically at interfaces, two possible mechanisms can be considered. In scenario (i), once the interface has become saturated, rodlet formation proceeds independently of the bulk phase and is governed exclusively by lateral interactions among RolA molecules already present within the interfacial film. Under this scenario, the lag time should scale with *C*_eq_, rather than with *C*_0_. Because *C*_eq_ is approximately independent of *C*_0_, no appreciable dependence of the lag time on the initial bulk concentration would be expected (Figure 6A). In contrast to this prediction, the experimentally determined lag time exhibited a clear dependence on *C*_0_ (Figure 3C). This result indicates that rodlet formation cannot be described as an ideal, bulk-independent reaction occurring merely within the pre-existing interfacial film. Instead, it supports scenario (ii), in which additional RolA molecules are continuously supplied from the bulk phase and adsorb onto and interact with the RolA film formed during the first plateau, thereby initiating and promoting rodlet formation (Figure 6B). This model is further supported by AFM images obtained during rodlet formation, which revealed the development of a bilayer-like RolA structure (Figures 2D, 3D). Consistent with these observations, previous structural analysis of RolA Langmuir–Blodgett film has also suggested that RolA forms film structures consisting of at least two molecular layers at the air–water interface [21]. Taken together, these findings suggest that rodlet formation involves not only lateral assembly within the initially adsorbed RolA layer but also additional adsorption and intermolecular interactions involving RolA molecules supplied from the bulk phase.

**Figure 6.**
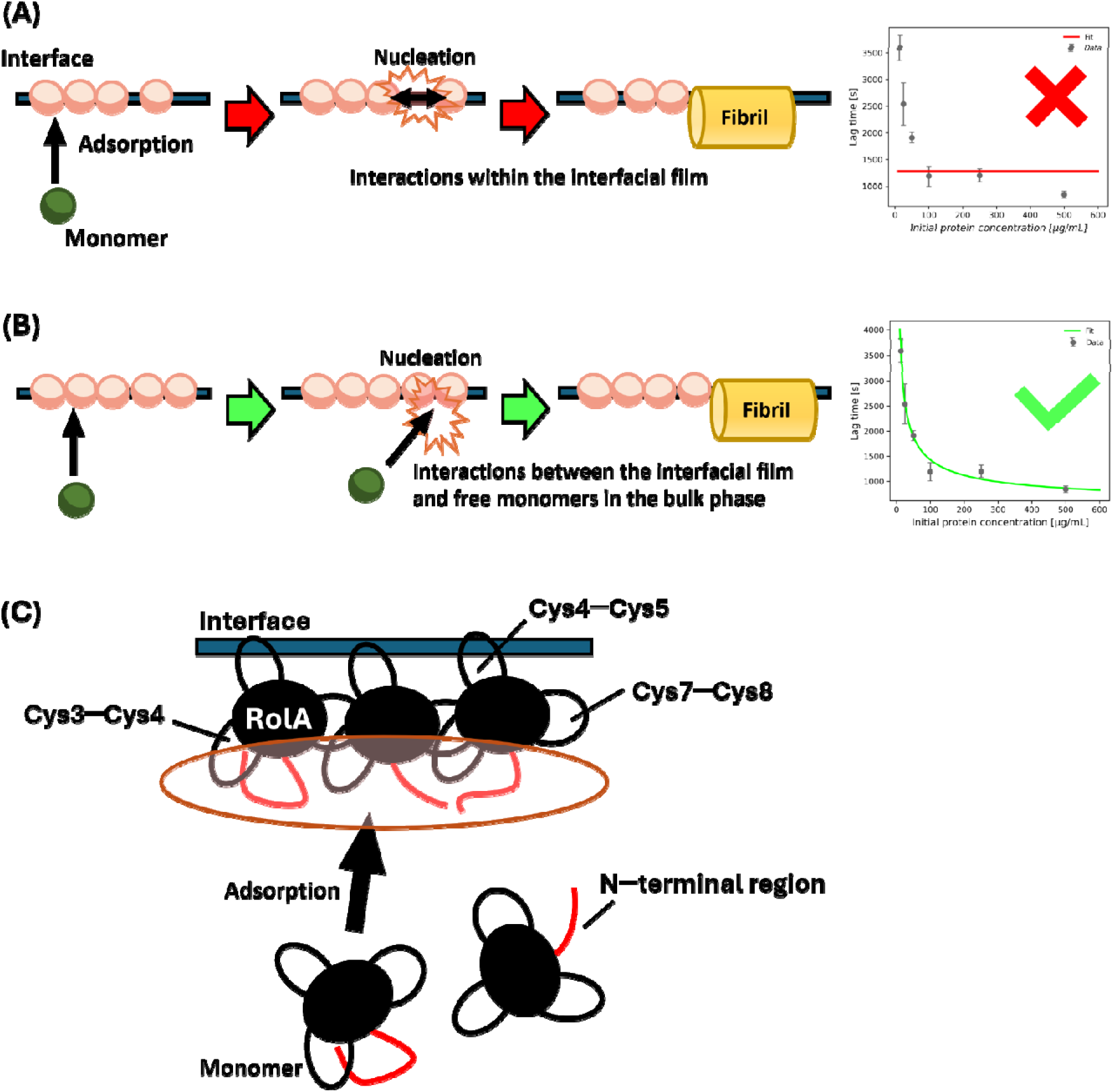
Possible models of RolA rodlet formation at the interface. (A) A model in which nucleation originates only from RolA adsorbed at the interface does not explain the experimentally observed dependence of lag time on the initial protein concentration. In contrast, the experimental data are consistent with (B) a model in which additional RolA monomers are supplied from the bulk phase to the interfacial RolA layer. (C) In the interfacial RolA film, the N-terminus tends to be oriented toward the aqueous phase and forms a catalytic field beneath the film. RolA monomers supplied from the aqueous phase to this surface may undergo surface-catalyzed nucleation in the vicinity of the film, leading to rodlet formation.

The difference in this interface-specific rodlet formation behavior among the RolA-WT and mutants suggests that the N-terminal region may play an important role at the interface. In the N-terminal mutants, the lag time in dynamic surface tension measurements was prolonged, indicating that nucleation was suppressed. Furthermore, the disappearance of this delay with addition of NaCl, which suppresses electrostatic interactions, suggests that electrostatic interactions were involved in nucleation. In the mutants used in this study, the N-terminal charged His and Lys residues were replaced with uncharged Ser residues, thereby altering the net charge of RolA and making it reasonable to expect changes in electrostatic intermolecular interactions. However, it remains unclear whether these mutations directly changed a specific function of the N-terminal region, and several possible scenarios need to be considered. For example, although RolA-H32S and -K34S have similar isoelectric points, they showed markedly different rodlet formation rates (Figures 2C, 4C), suggesting that the complex mechanism of RolA rodlet formation cannot be explained simply by net charge. The detailed mechanism underlying these differences should be clarified.

To begin to address this knowledge gap, we focused on the interaction between the interfacial RolA film and RolA in the bulk phase as mediated by the N-terminus. To grasp a molecular-level picture of the role of the N-terminal region, MD simulations were performed and revealed that the aqueous-phase side of interfacial RolA film is composed of the Cys3– Cys4 loop and the N-terminal region. Because rodlet formation is driven by the approach of additional RolA molecules from the bulk phase to the interfacial RolA film (Figure 6B), the aqueous-facing region of interfacial RolA may be involved in the surface-catalyzed rodlet formation occurring on the RolA film (Figure 6C). The N-terminal IDR region may mediate intermolecular interactions through its high mobility. However, when the positive charge is reduced by mutagenesis, intramolecular interactions within the N-terminal region may become dominant (Figure 5D). As a result, opportunities for intermolecular interactions mediated by the N-terminal region may be reduced, which could lead to a slower rate of rodlet formation. Another possibility that cannot be excluded is that the negative charge of the Cys3–Cys4 loop (residues 94–101, GLGLLDEC), which tends to form transient contacts with the second half of the N-terminal region in WT, may actually have an inhibitory effect on rodlet formation due to its charge. In this case, the N-terminal region may facilitate rodlet formation in WT by masking these negative charges, whereas their exposure in H32S/K34S may reduce the rodlet formation propensity (Figure 5E). From a macroscopic mean-field perspective, changes in intramolecular interactions involving the N-terminal region alter the properties of the aqueous-facing surface of the interfacial film, thereby modulating the rate of surface-catalyzed rodlet formation. This interpretation is consistent with heterogeneous nucleation at interfaces, in which a surface promotes nucleation by lowering the free-energy barrier for formation of a critical amyloid nucleus relative to homogeneous nucleation in bulk solution [29].

Given their structural similarity to common amyloid fibrils, hydrophobin rodlets can reasonably be compared to others within the wider category of amyloid fibrils. Surface-catalyzed effects have been reported in diverse amyloid systems [29,30], and the mechanism described in this study may represent an example of this broader phenomenon. In well-established amyloid systems, for example, fibril surfaces promote nucleation and can cause a rapid increase in fibril mass through secondary nucleation [31]; surface-catalyzed elongation that facilitates fibril alignment has also been reported recently [26]. In addition, the air–water interface [30], solid–liquid interface [22,32], lipid membrane surface [33], and the surfaces of protein condensates formed by liquid–liquid phase separation (i.e., liquid–liquid interfaces) [34,35] have all been shown to express catalytic effects that either promote or suppress fibril formation. The system identified in this study—rodlet formation promoted via catalytic effect at the surface of an amorphous RolA film—extends the conceptual view of surface-catalyzed reaction in amyloid fibrillation. Amyloid fibril formation is often regarded as analogous to crystal growth [36]. In crystal growth, the speed and polymorphism of crystal growth beneath surfactant monolayers at the air– water interface depends on the surface charge state and the types of functional groups exposed at the monolayer surface [37]. Therefore, rodlet formation beneath the interfacial RolA monolayer may be interpreted as analogous to such monolayer- templated crystal growth.

The utilization of the N-terminal region of RolA is also unique from a biological perspective. RolA adsorbs onto solid polymers, coats the surface, and subsequently recruits polymer-degrading enzymes via its N-terminal region, thereby enhancing substrate degradation [8,9]. Moreover, the present study shows that RolA controls its rodlet formation through the N-terminal region (Figure 6C). Thus, the N-terminal region of RolA can be regarded as a bifunctional segment. IDRs can modulate their structural states and functional outputs in response to interaction partners owing to their conformational flexibility [38]. This moonlighting behavior could enable RolA to simultaneously assume diverse roles and to function as a hub that coordinates complex biological processes. Because the relationship between the diverse biological functions reported for various hydrophobins and their N-terminal regions remains unclear in almost all cases, further detailed analyses are required in future studies.

## CONCLUSION

In this study, the interface-specific mechanism of RolA rodlet formation and the influence of its N-terminal region on rodlet formation were elucidated through kinetic analyses based on AFM observation, dynamic surface tension measurements, and MD simulations. Initially, RolA formed an amorphous film at the interface, thereby establishing a reaction field for rodlet formation. Then, rodlets were formed by interacting with this film, with a portion of the Cys3–Cys4 loop and N-terminus exposed on its surface. These findings highlight the potential of the RolA N-terminal region to mediate diverse functions and provide a conceptual advance by updating the models for heterogeneous and complex pathways of interfacial rodlet formation.

## MATERIALS & METHODS

### Protein expression and purification

The *A. oryzae* RolA WT-overexpressing strain, *A. oryzae* RolA H32S-overexpressing strain, *A. oryzae* RolA K34S- overexpressing strain, and *A. oryzae* RolA H32S/K34S-overexpressing strain [9] were grown in YPM liquid medium (1% yeast extract, 2% polypeptone, and 2% maltose). Conidiospores of *A. oryzae* were inoculated into YPM liquid medium at 1 × 10^6^ spores/mL and cultivated with shaking at 30 °C for 48 h (WT), 34 h (H32S and K34S), or 24 h (H32S/K34S). The culture was filtered through Miracloth (Merck kGaA, Darmstadt, Germany), and the obtained culture filtrate was adjusted to pH 8.5 with 100 mM Tris (pH 10.5), and then the electrical conductivity was adjusted to about 1.00 with Milli-Q water (MQ). The solution was subjected to a Cellufine Q-500 column (JNC Corp., Tokyo, Japan) buffered with 10 mM Tris-HCl (pH 9.0). RolA was eluted with a 0.05–0.3 M linear gradient of NaCl. Eluted fractions containing RolA were dialyzed against 10 mM sodium-citrate buffer (pH 4.0) and then submitted to SP-Sepharose fast flow column (GE Healthcare, Tokyo, Japan) buffered with the same buffer. RolA was eluted with a 0–0.3 M linear gradient of NaCl. The purified RolA was dialyzed against 10 mM ammonium acetate (pH 7.0), then lyophilized.

### Preparation of hydrophobic substrate

Silicon wafers (p-Si wafers, ≤0.02 Ω cm; Mitsubishi Materials Trading Co., Tokyo, Japan) were washed via ultrasonication in chloroform, acetone, and 2-propanol for 15 min respectively, and were dried with N_2_ gas. Then, the surface of the wafers was cleaned with an ultraviolet ozone cleaner (SKB401Y-01; SUN ENERGY Co. Ltd., Kanagawa, Japan) for 30 min. Finally, the wafers were immersed in chloroform solution containing 0.1% n-octyltrichlorosilane (Tokyo Chemical Industry Co. Ltd., Tokyo, Japan) and incubated overnight to make the surface hydrophobic.

### Atomic force microscopy observation

Rodlet formation speed assessment was performed according to Takahashi et al. [19] with modifications (Figure 2A). Pedestals were placed in septum-sealed vials (1114-S004; Quality Environmental Containers, Beaver, USA) containing 10 mM sodium-acetate buffer (pH 5), and Parafilm was placed on top of each pedestal. The vials were sealed and incubated until the humidity exceeded 90%, and the humidity inside the vial was monitored with a humidity indicator card (RH3-70; As One, Osaka, Japan). Then, 20 µL of purified RolA solution (100 µg/mL; 7.35 μM) prepared with 10 mM sodium-acetate buffer (pH 5) was injected into the vials using a syringe (water-in-air sessile droplet). All procedures were conducted at 23 °C. After the injection, droplets of RolA solution were incubated for 10–240 min at 23 °C. After the incubation, the vial cap was gently opened and a hydrophobic substrate was pressed against the droplet surface for 30 s. The substrate was then immersed in a large volume of MQ to remove excess protein solution and allowed to dry for at least 1 h.

AFM observations were performed using a scanning microscope (SPA400; Seiko Instruments Inc., Chiba, Japan) and a cantilever (SI-DF20, Al-coated; f = 136 kHz, C = 16 N/m; Hitachi High-Tech Science Corp., Tokyo, Japan). Topographic images were acquired in dynamic surface mode over a scan area of 1000 × 1000 nm. For each time point, one topographic image of the RolA-transferred film was obtained from each of three independent experiments. The height and length of rodlets were measured using ImageJ 1.54d (http://imagej.org).

### Dynamic surface tension measurement

Dynamic surface tension was measured by the pendant drop method [39] using a Drop Master 300 contact angle meter (Kyowa Interface Science Co., Ltd., Saitama, Japan). Immediately before measurement, lyophilized RolA was dissolved in 10 mM sodium-acetate buffer (pH 5) and passed through a filter (0.22 µm; Hawach Scientific, Shaanxi, China). The solution was centrifuged at 20,000 × *g* for 30 min at 4 °C, and 80% of the supernatant was collected. The RolA concentration was determined using a spectrophotometer (V-670; JASCO, Tokyo, Japan) and adjusted to the desired concentration by dilution. Then, RolA solution was dispensed from the tip of a syringe into a sealed chamber to form a 7.5- µL droplet and incubated for 7200 s at room temperature (23 °C). To minimize the evaporation of the droplet during the measurement, 10 mM sodium-acetate buffer (pH 5) was dispensed into the chamber and incubated until the humidity exceeded 90%. During measurements, to prevent excessive pressure buildup inside the chamber and backflow of the solution into the syringe, a needle was inserted into the Parafilm to create a very small vent. Droplet images were captured every 30 s for 2 h with a CCD camera, and the surface tension of the solution was calculated using FAMAS software (Kyowa Interface Science Co., Ltd., Saitama, Japan), in which the droplet shape line was fitted to the Young–Laplace equation to determine surface tension. Three independent measurements were performed for each RolA concentration.

The relationship between surface tension and the interfacial concentration of the protein can be described by the Gibbs adsorption isotherm as follows:

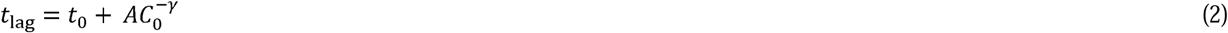

where *R* is the gas constant, *T* is the absolute temperature in K, γ is the interfacial tension, and *C* is the RolA concentration. Using this equation, *Γ* was determined from the slope of the linear fit to the plot. The concentration at which the surface tension reached a minimum in the linear fit was defined as the CMC.

### MD simulation conditions

All-atom MD simulations were performed with the CHARMM36m force field [40] using LAMMPS software [41]. For both the RolA-WT and -H32S/K34S structural model, two types of simulations were performed: one in bulk solution and the other at the interface. The RolA structure was predicted using Alphafold2 [42] (Figure 1) and used as the WT model (RolA- WT). Based on this structure, His32 and Lys34 were replaced with Ser, and the resulting model was used as the RolA- H32S/K34S model. In all simulation systems, the temperature was maintained with a Nosé–Hoover thermostat [43,44], and the pressure was controlled with a Parrinello–Rahman barostat [45,46]. The equations of motion were integrated using the Verlet algorithm [47] with a time step of 2 fs, along with the SHAKE algorithm [48], to constrain the bond lengths to hydrogen. The nonbonded interactions were calculated with a cutoff distance of 1.2 nm, and the particle–particle–particle– mesh (PPPM) method [49] was used to calculate long-range electrostatic interactions. For the simulations in the bulk phase, RolA-WT and -H32S/K34S structural models were placed in each simulation box (71 × 71 × 71 Å) and solvated by adding water. After the steepest-descent energy minimization, the systems were relaxed for 50 ps at a temperature of 300 K and a pressure of 1 atm, followed by production runs of 200 ns in the NPT ensemble at a temperature of 310 K under three-dimensional periodic boundary conditions. For the simulations at the interface, a 71 × 71 × 70 Å water slab was placed in the center of a 71 × 71 × 106 Å simulation box, thereby creating air (vacuum) to water interfaces. The final frames of the bulk simulations of RolA-WT and -H32S/K34S were used as the initial structures and placed in the water slab (Figure 5A). Production simulations of RolA adsorption at the air–water interface were performed for 200 ns in the NVT ensemble.

For WT adsorption at the interface, four independent simulations starting from different initial orientations were conducted. Here, we checked the time evolution of the *z*-coordinates of the centers of mass calculated using the C_α_ atoms of residues 76–86, 104–115, and 132–143, corresponding to the Cys3–Cys4, Cys4–Cys5, and Cys7–Cys8 loops, respectively. In all four simulations, the Cys4–Cys5 loop was oriented toward the air phase, whereas the Cys3–Cys4 loop was oriented toward the aqueous phase (Figure S5). Since RolA adopted a stable orientation at the interface within 80 ns in all four simulations, we confirmed that 100–200-ns trajectories were suitable for analyzing RolA structure in its stable interfacial state. The probability density distributions were calculated over 100–200 ns using the C_α_ coordinates of the three loop regions described above and the N-terminal region (residues 17–51) (Figure 5B). In the H32S/K34S simulations, the interfacial structure adopted an orientation similar to that of WT within 80 ns, with the Cys4–Cys5 loop oriented toward the air phase and the Cys3–Cys4 loop oriented toward the aqueous phase (Figure S6), so we confirmed that 100–200-ns trajectories were suitable for performing structural analysis. To analyze the intramolecular contacts of the N-terminal region with other regions, contact probabilities were calculated separately for the first (residues 18–31) and second (residues 32– 44) halves of the N-terminal region. Two residues were considered to be in contact in each frame when at least one pair of their heavy atoms was within less than 4.5 Å [50]. The contact probability for each residue pair was defined as the number of frames in which the contact occurred divided by the total number of analyzed frames, using trajectory data for 100–200 ns. To generate one-dimensional contact profiles, the contact probabilities between each target residue and all residues within each half of the N-terminus were averaged. MDAnalysis [51,52] was used to calculate the RMSF of the C_α_ atoms over the 100–200 ns of each trajectory relative to the average structure obtained from the trajectory. VMD (ver. 1.9.4a53) was used to generate the image.

## AUTHOR CONTRIBUTIONS

Conceptualization, N.T., and K.A.; Formal analysis, N.T. and N.A.; Methodology, N.T., N.A., T.M., M.F., H.Y., and K.A.; Writing-original draft, N.T., N.A., Y.T., T.T., A.Y., and K.A.; Writing-review and editing, T.M., M.F., and H.Y.; Funding acquisition, N.T., and K.A.

## ACKNOWLEDGMENTS

This work was supported by the Japan Society for the Promotion of Science under a Grant-in-Aid for Scientific Research (B) [23K26810] to K.A. and a Grant-in-Aid for JSPS Fellows [24KJ0419] to N.T., and received funding from the Noda Institute for Scientific Research (K.A.).

