## Supporting Information for "N-terminal intrinsically disordered region mediates self-catalytic interfacial nucleation of *Aspergillus oryzae* hydrophobin RolA"

Figure S1

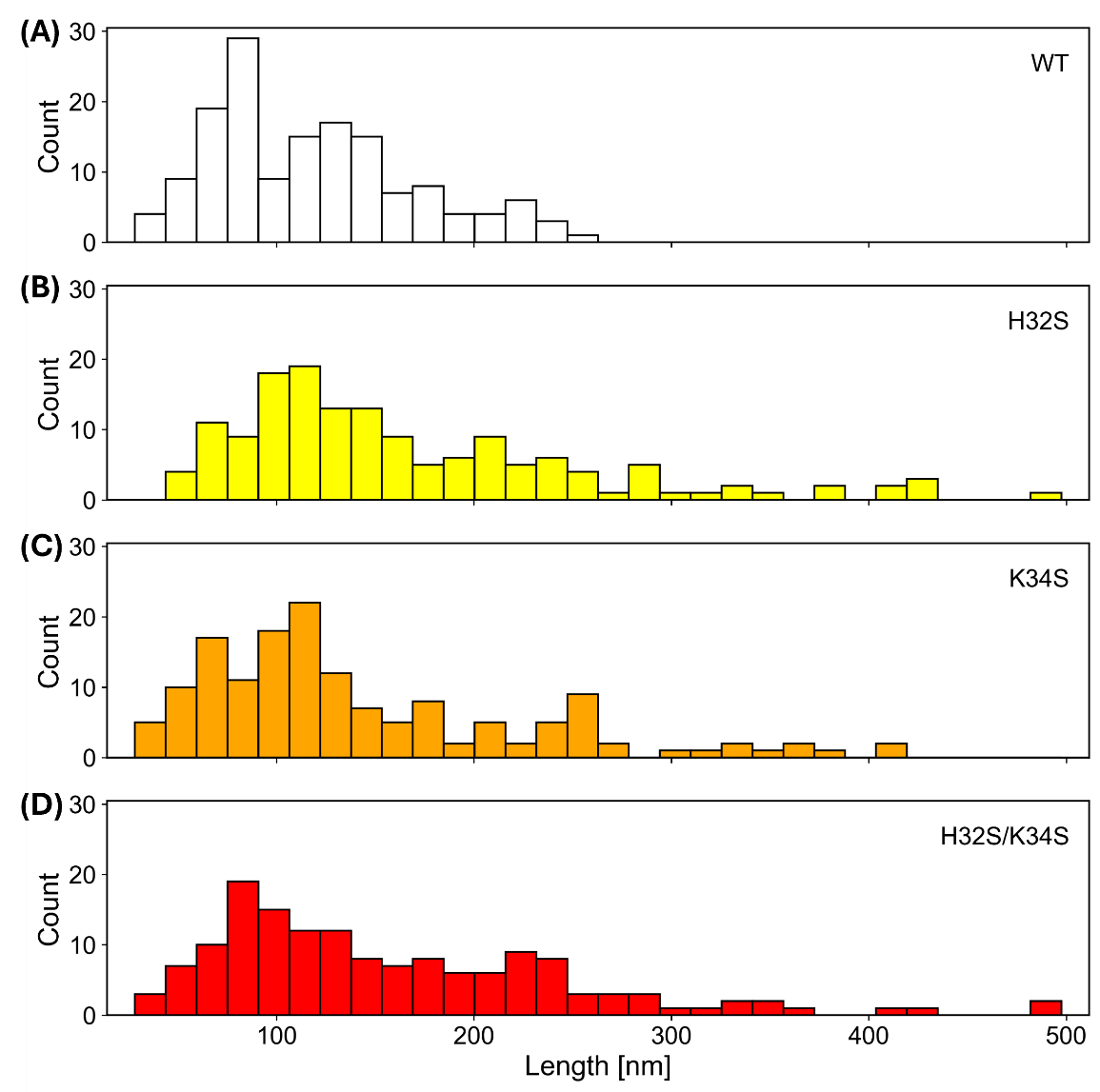

Length distributions of rodlets formed on the surface of sessile water droplets for (A) RolA-WT, (B), -H32S, (C) K34S, and (D) -H32S/K34S.

Figure S2

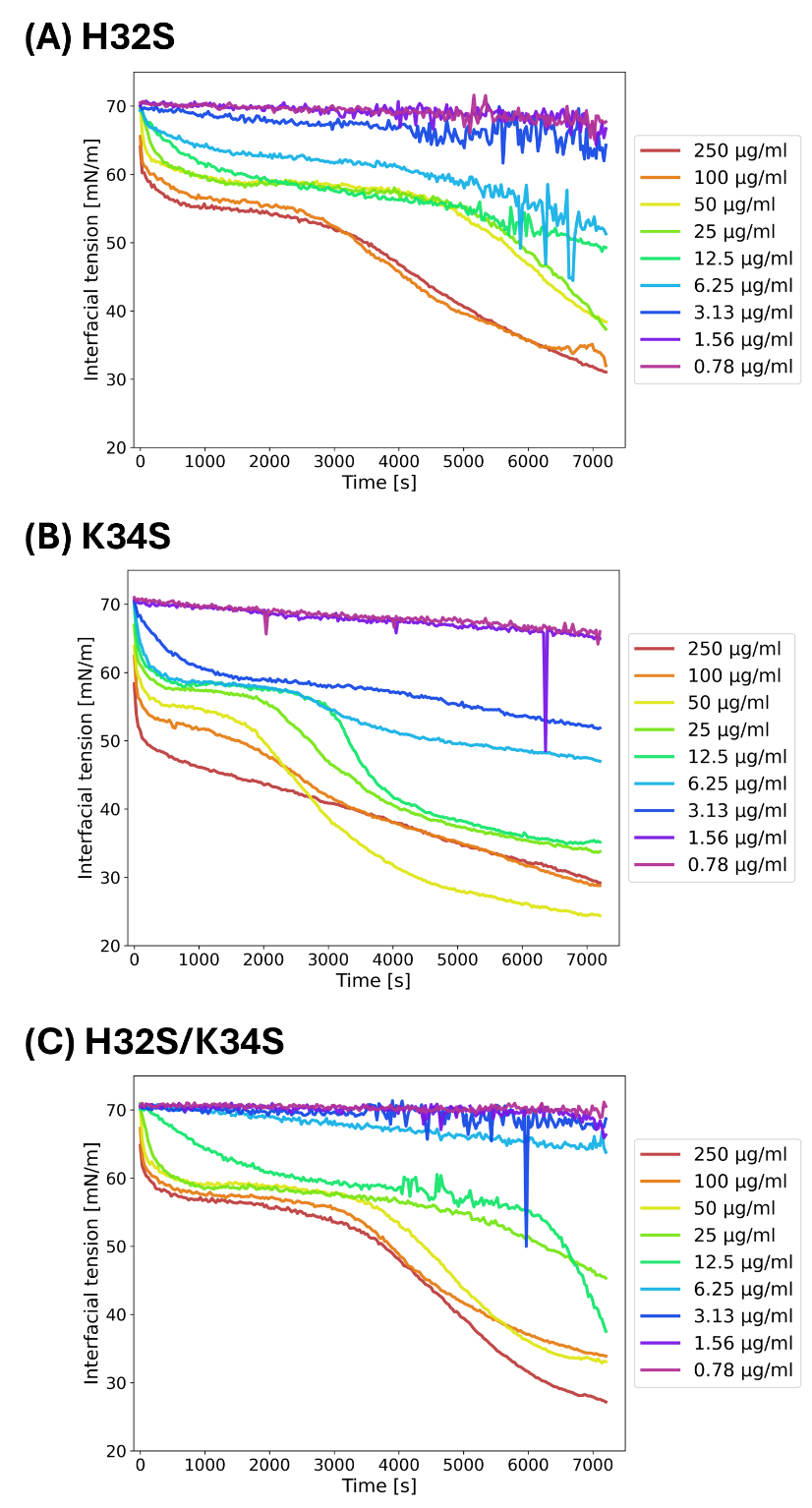

Dynamic surface tension profiles of (A) -H32S, (B) -K34S, and (C) -H32S/K34S at various initial concentrations.

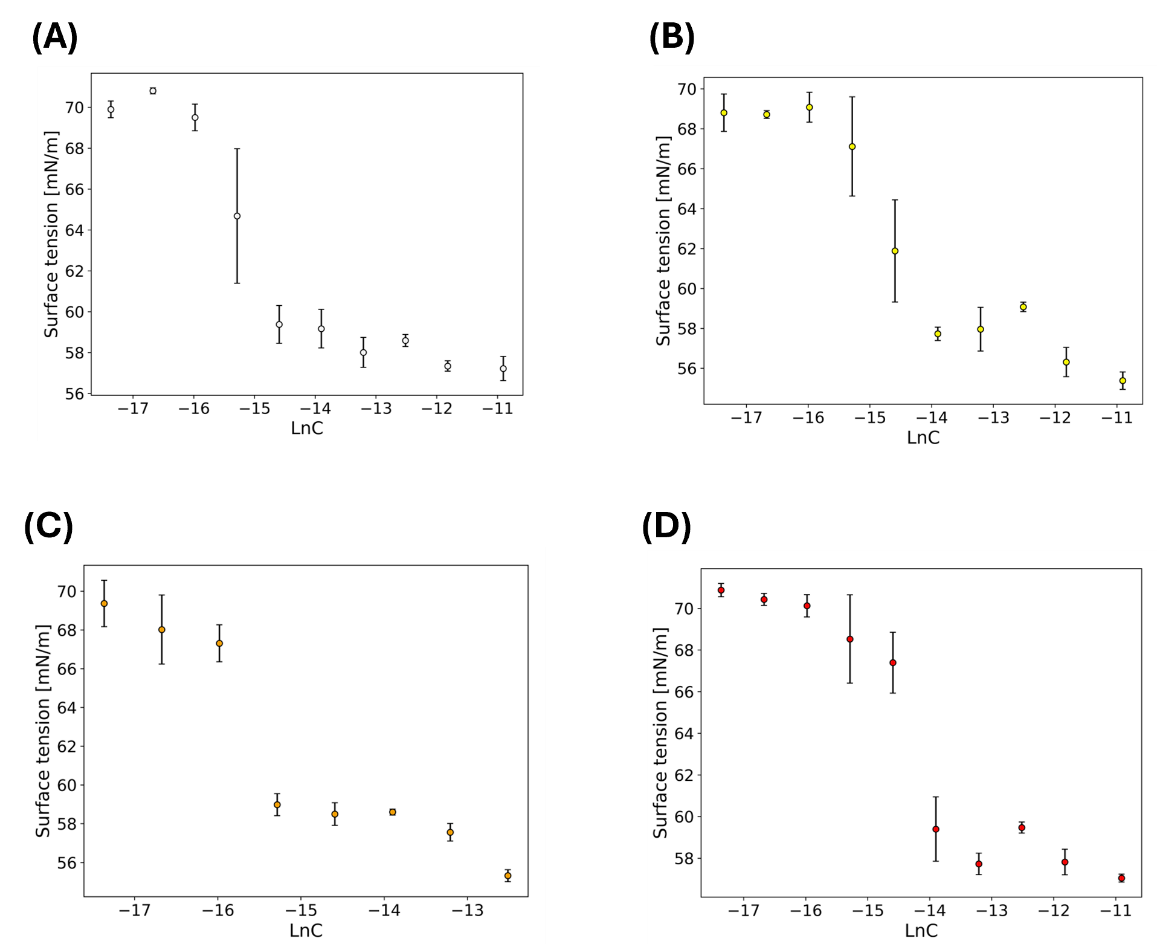
Figure S3

Equilibrium surface tension vs. initial protein concentration (lnC) for (A) RolA-WT, (B), -H32S, (C) -K34S, and (D) -H32S/K34S. Error bars show standard deviation (*n* = 3).

Figure S4

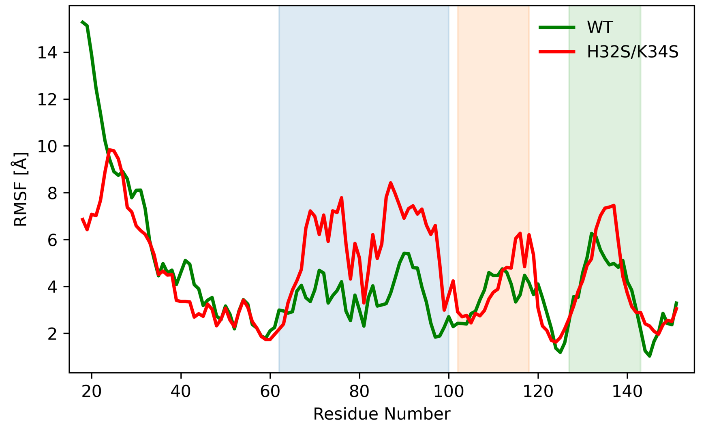

Root-mean-square fluctuation (RMSF) of the C_α_ atoms of each residue for RolA-WT (green) and -H32S/K34S (red). The Cys3–Cys4, Cys4–Cys5, and Cys7–Cys8 loops are indicated by blue, orange, and green shading, respectively.

Figure S5

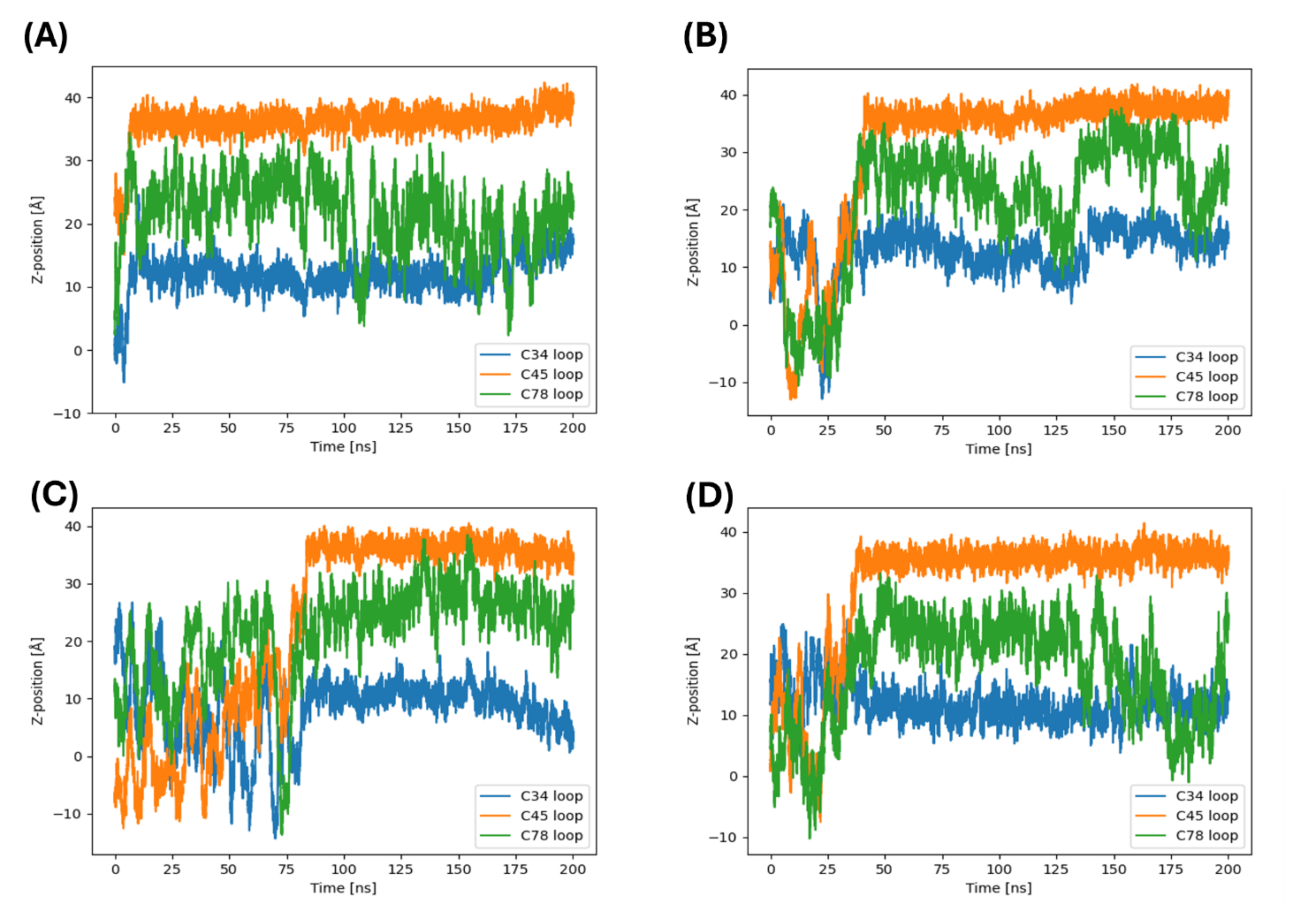

Time evolution of *z*-positions of the Cys loops in RolA-WT. The four panels show the results of four independent simulations with different initial orientations.

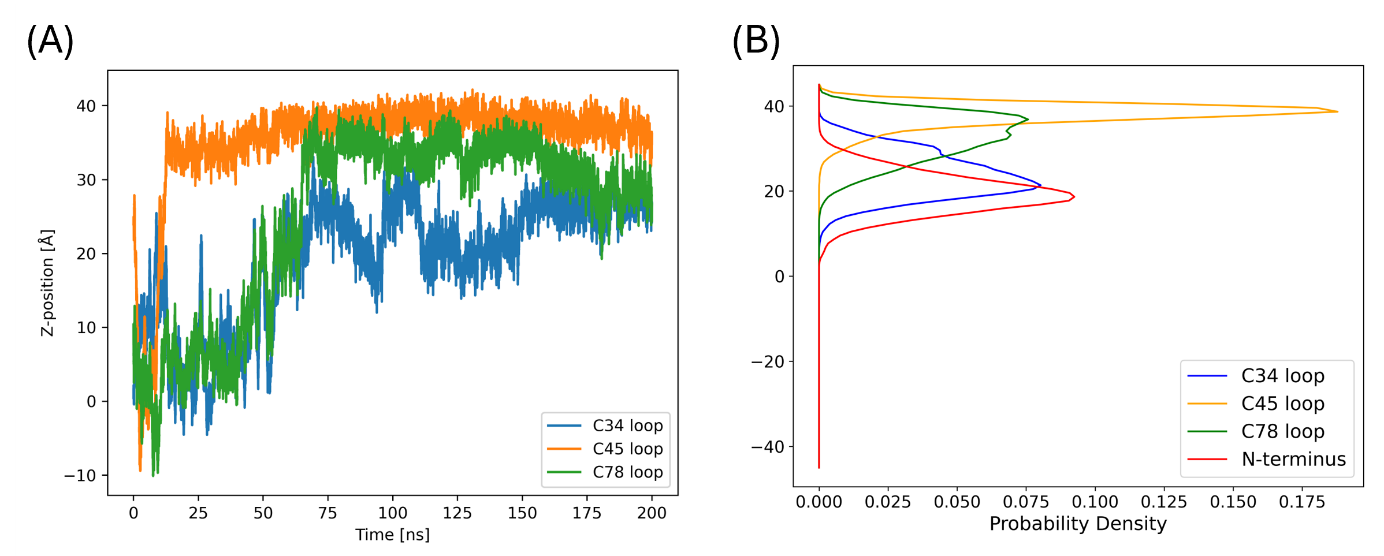
Figure S6

Orientation of RolA-H32S/K34S at the air–water interface. (A) Time evolution of the *z*-positions of the Cys loops. (B) Spatial distribution of the Cys loops and the N-terminal region.

Table S1. Surface active properties of RolA-WT and three mutants.

|  | *Γ*_max_ (×10^−6^) (mol/m^2^) | CMC (μg/mL) |
| --- | --- | --- |
| WT | 2.96 ± 0.08 | 6.25 |
| H32S | 2.74 ± 0.63 | 12.5 |
| K34S | 2.16 ± 0.29 | 6.25 |
| H32S/K34S | 2.36 ± 0.61 | 25 |

*Γ*_max_, maximum surface excess concentration; CMC, critical micelle concentration.
